# Antibody-mediated rescue of endogenous retrovirus-induced damage in the demyelinated central nervous system

**DOI:** 10.64898/2026.06.10.731326

**Authors:** Laura Reiche, Joel Gruchot, Benjamin Charvet, Hans-Peter Hartung, Margot Lemarinier, Alexandre Lucas, Hervé Perron, David Leppert, Celine Heeb, Urs Meyer, Patrick Küry

## Abstract

The human endogenous retrovirus type W (HERV-W) has been identified as a human-specific neuropathological factor that preferentially affects glial cell types in multiple sclerosis (MS). Recent work using transgenic mice with expression of the HERV-W envelope (ENV) protein unveiled that this endogenous retroviral element disrupts myelin repair and polarizes microglial and astroglial cells towards axon-damaging neurotoxic phenotypes. Moreover, initial clinical trials using Temelimab, a neutralizing antibody targeting the HERV-W ENV protein, have provided circumstantial evidence that ENV exerts anti-regenerative and neurodegenerative effects in MS patients. Aligning these observations, it was therefore concluded that HERV-W represents an important factor contributing to disease progression independent of relapse activity (PIRA). Building on these findings, we here applied a neutralizing anti-ENV antibody in a non-inflammatory demyelination mouse model to directly evaluate its potential to mitigate neurodegeneration and ameliorate remyelination. In transgenic mice with human-specific expression of the HERV-W ENV protein, repetitive intraperitoneal anti-ENV antibody injections resulted in accelerated oligodendroglial differentiation, enhanced remyelination, axonal protection, and reduced neurofilament light chain leakage in the serum. Neurotoxic microglial traits were also reduced, while homeostatic parameters were stabilized. As astroglial cells underwent a similar shift, inducing regenerative traits at the expense of toxic parameters, anti-ENV application overall generated a less hostile cellular environment. This study provides direct evidence of the capacity of HERV-W neutralizing antibodies to access the central nervous system and to ameliorate damage conferred by this viral entity previously associated with smouldering disease processes.

**Significance Statement:** Although neurodegeneration is a hallmark of multiple sclerosis (MS), its underlying mechanisms are poorly understood. Clinically, it manifests as smouldering MS or progression without relapse activity (PIRA). This is the primary factor leading to the accumulation of clinical disability and is currently untreatable. This study provides the first direct evidence that antibodies directed against the HERV-W ENV protein can attenuate the activity of neurodegeneration-promoting glial cells *in vivo*. Our data validates neutralization of this endogenous retroviral element as a promising therapeutic approach particularly relevant to the chronic form of MS.

## Introduction

The identification of disease-conferring factors constitutes a first critical step and is crucial when addressing complex neurological diseases such as multiple sclerosis (MS). In contrast to the almost complete suppression of relapse activity by high-efficacy therapies such as anti-CD20-antibodies (1), the mode of action of candidate compounds trying to attenuate progression independent of relapse activity (PIRA) were largely unsuccessful in clinical trials (2). However, additional challenges arise from the design of potential counteracting measures and their evaluation in terms of central nervous system (CNS) penetrance and effectiveness. Furthermore, as MS is caused by overlapping pathological processes and involvement of multiple cell populations, the identification of targets for therapeutic intervention is complicated.

As immune-mediated demyelinating disease, MS is primarily characterized by peripheral immune cell infiltration, focal inflammation resulting in lesion formation in white and grey matter and the loss of oligodendrocytes and myelin sheaths, ultimately leading to neuronal death. PIRA phases, on the other hand, show compartmentalized, low-grade inflammation with limited peripheral immune cell infiltration while brain-diffuse neurodegenerative processes predominate primarily due to neurotoxic microglia and astrocytes (3, 4).

The human endogenous retrovirus type-W (HERV-W) constitutes dormant retroviral sequences embedded in the human genome that can specifically be activated in a few pathological situations, such as in MS (see for reviews (5, 6). While transcript and some HERV-W encoded proteins were found to gradually increase in expression during disease evolution and progression (7, 8), numerous studies demonstrated that the encoded envelope (ENV) protein is primarily responsible for its pathological impact.

In this context, initial investigations focused on the immune response to HERV-W ENV (9-11). Subsequently, a substantial negative impact of the ENV protein on the myelin repair potential (12) and concurrent neurodegenerative traits were described (13). Consequently, converging evidence emerged on specific HERV-W ENV-mediated effects towards oligodendrocytes, microglia and astrocytes (14).

Due to its human-specific origin, knowledge of HERV-W’s role towards glial cells has evolved through a combination of various observations. These include histological assessments of MS-derived brain tissue which were subsequently corroborated functionally by *ex vivo* cell culture studies (12, 13). A detailed examination of a novel transgenic mouse line that mimicked endogenous HERV-W ENV expression in the diseased CNS provided additional *in vivo* evidence of its functionality (14). Finally, indirect evidence of its implication in human neurodegeneration and white matter repair surfaced in two clinical trials using a HERV-W ENV protein-neutralizing antibody, Temelimab. In participants with MS that received this antibody, brain atrophy levels were reduced, whereas signs of improved myelin integrity and stability were observed (15).

The investigation presented here sought to bridge these observations and to further strengthen the evidence on the detrimental role of HERV-W in MS. Furthermore, it aimed to demonstrate the deposition of neutralizing antibodies within affected tissues substantiating the therapeutic value of Temelimab. To this end, the HERV-W ENV protein-neutralizing antibody was injected into the demyelinating and degenerating transgenic mice with human-specific expression of HERV-W ENV and found to penetrate the brain parenchyma and to rescue tissue damage related to axon myelination and axonal integrity.

## Results

### Anti-ENV application in demyelinating transgenic HERV-W ENV mice rescues myelin content and reduces axonal degeneration

As HERV-W constitutes a human-specific pathogenic entity, we previously assessed the impact of the HERV-W ENV protein in a functional *in vivo* context using a specific transgenic mouse model (14). This mouse exhibits a moderate expression of HERV-W ENV, comparable to levels observed in human tissue (7, 16). A detailed analysis of these mice revealed significant deficits in myelin repair following demyelination induced by cuprizone treatment. This was accompanied by the presence of neurotoxic microglia and astroglia. Furthermore, HERV-W ENV mice exhibited substantially heightened signs of axon degeneration. Moreover, following immunization, experimental autoimmune encephalomyelitis (EAE) activity progressed faster in transgenic mice, accompanied by activated glial cells. This study, therefore, provided the first direct evidence of HERV-W ENV’s contribution to the overall negative impact of this activated viral entity in MS-related neuropathology.

Using the same cuprizone-mediated demyelination model as in our previous study (14), we here tested the ability of an intraperitoneally (i.p.) injected anti-ENV antibody to ameliorate HERV-W-specific pathological changes in the transgenic mouse line. For maximum comparability and immune compatibility, we used the original mouse anti-ENV antibody (ENV01, GeNeuro) which was subsequently humanized to generate the clinically administered anti-ENV antibody Temelimab (15). Cuprizone-mediated demyelination of the corpus callosum was selected over EAE as we specifically wanted to address non-immune cell-dependent effects. Additionally, given the critical role of HERV-W in smouldering disease processes (14, 17), a mostly intact blood-brain-barrier (BBB) was anticipated. Consequently, pathological processes within the parenchyma were of particular interest for the present study. Transgenic mice (hemizygote males and homozygote females) were fed a diet containing 0.2% cuprizone (CPZ) for 7 weeks. Subsequently, they were switched to a control diet to induce myelin repair. During cuprizone-mediated demyelination of transgenic mice, either the specific anti-ENV antibody or a matching isotype control antibody was injected three times i.p. (Fig. 1A). When comparing mice treated with anti-ENV versus control antibody, Luxol fast blue (LFB) staining of the caudal corpus callosum revealed rescued myelinated areas, both during de- and remyelination time-points (Fig. 1B,D). Notably, there was a marked reduction in the number of amyloid precursor protein (APP)-positive spheroids (Fig. 1C,D), which serves as a surrogate marker for axon degeneration (14, 18).

**Figure 1:**
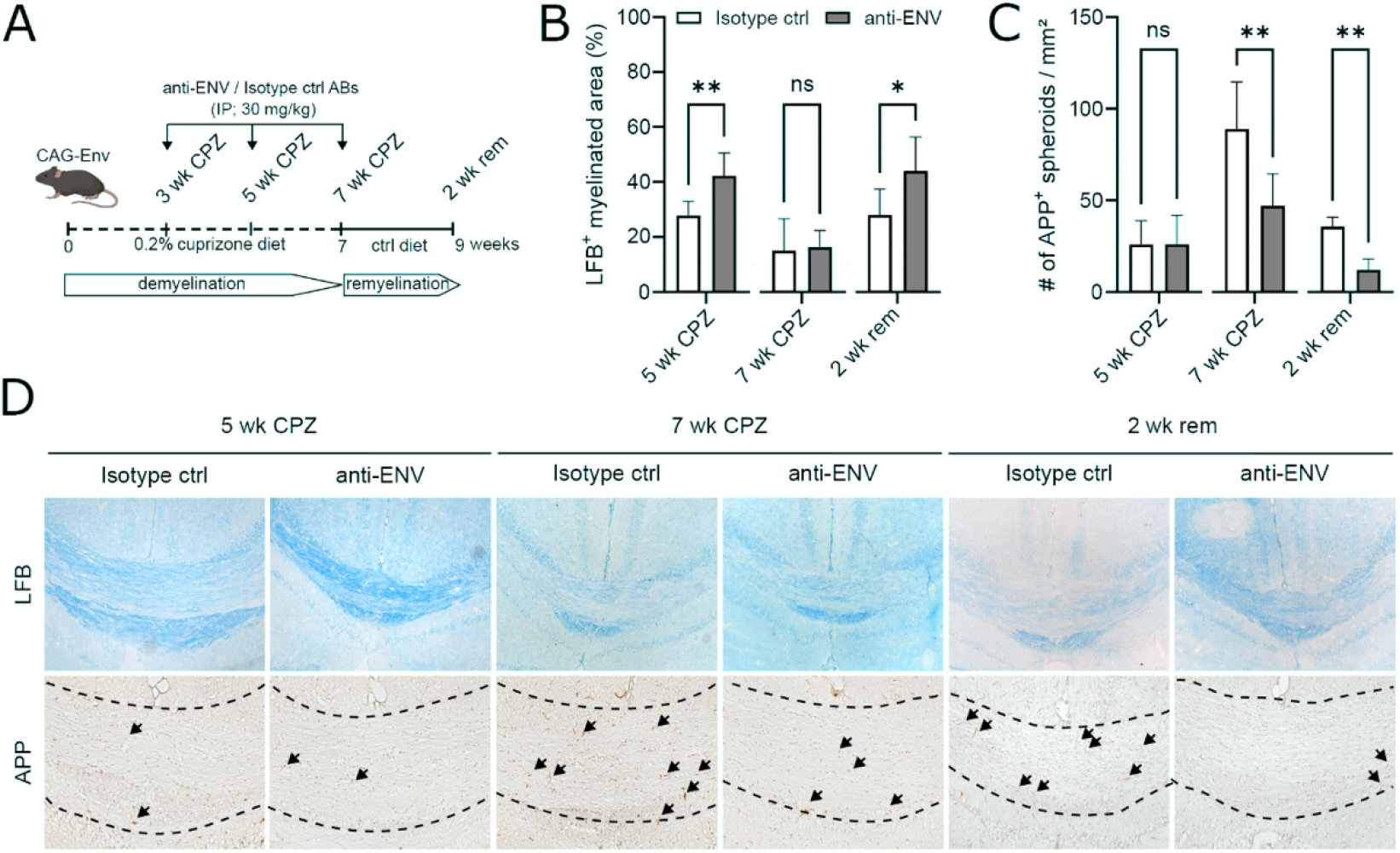
Anti-ENV administration ameliorates remyelination and reduces axonal degeneration in transgenic HERV-W ENV expressing mice. (A) Cuprizone (CPZ)-mediated demyelination of the corpus callosum and anti-ENV or isotype control antibody injections in HERV-W ENV expressing transgenic mice. (B) Anti-ENV-dependent rescue of myelinated areas. (C) Reduction in APP-positive spheroids marking damaged axons in anti-ENV treated mice. (D) Representative images of Luxol fast blue (LFB)- and anti-APP-stained tissue sections. Arrows point to APP-positive spheroids. Data are presented as mean values (B and C: n = 6) ± SEM. Statistical significance of was assessed by Student’s two-sided, unpaired t-test and Mann-Whitney *U* two-tailed test. Data were considered as statistically significant (95% CI) at *P < 0.05, **P < 0.01. n.s. = not significant. wk = week, ctrl = control. (Scale bar: 100 µm).

### Presence of the hexameric ENV protein and anti-ENV antibodies in mouse serum and brain parenchyma

We previously confirmed HERV-W ENV protein expression in our transgenic mouse model (14), however, Charvet and colleagues reported an hexameric extracellular form of the ENV protein in humans relevant for solubility and antibody recognition (19). We therefore applied improved detection protocols on transgenic mouse sera and brain extracts to further classify the ENV protein forms occurring in our transgenic mouse strain. Using automated Simple Western technology, we demonstrated that the ENV protein in transgenic mice appears to be primarily present as hexamers (Fig. 2A), similar to what has been described in human patient sera and, originally identified in brain samples from MS lesions (19). Anti-ENV antibody detection after two injections was then done by means of tissue serology (as outlined in Fig. 2B). This revealed that the anti-ENV antibody, specifically its HERV-W ENV protein binding activity, was only detectable in sera from transgenic mice injected with the antibody but not upon control antibody administration (Fig. 2C). More importantly and particularly relevant in the context of clinical applications, we confirmed the presence of the anti-ENV antibody in the perfused and non-fixed brain parenchyma of transgenic mice (Fig. 2D). Overall, these protein data unequivocally demonstrated that the ENV protein is present in its pathological appearance and that i.p. administration of the anti-ENV antibody in these transgenic mice results in detectable amounts of this antibody behind the intact BBB, thereby constituting a suitable therapeutic approach.

**Figure 2:**
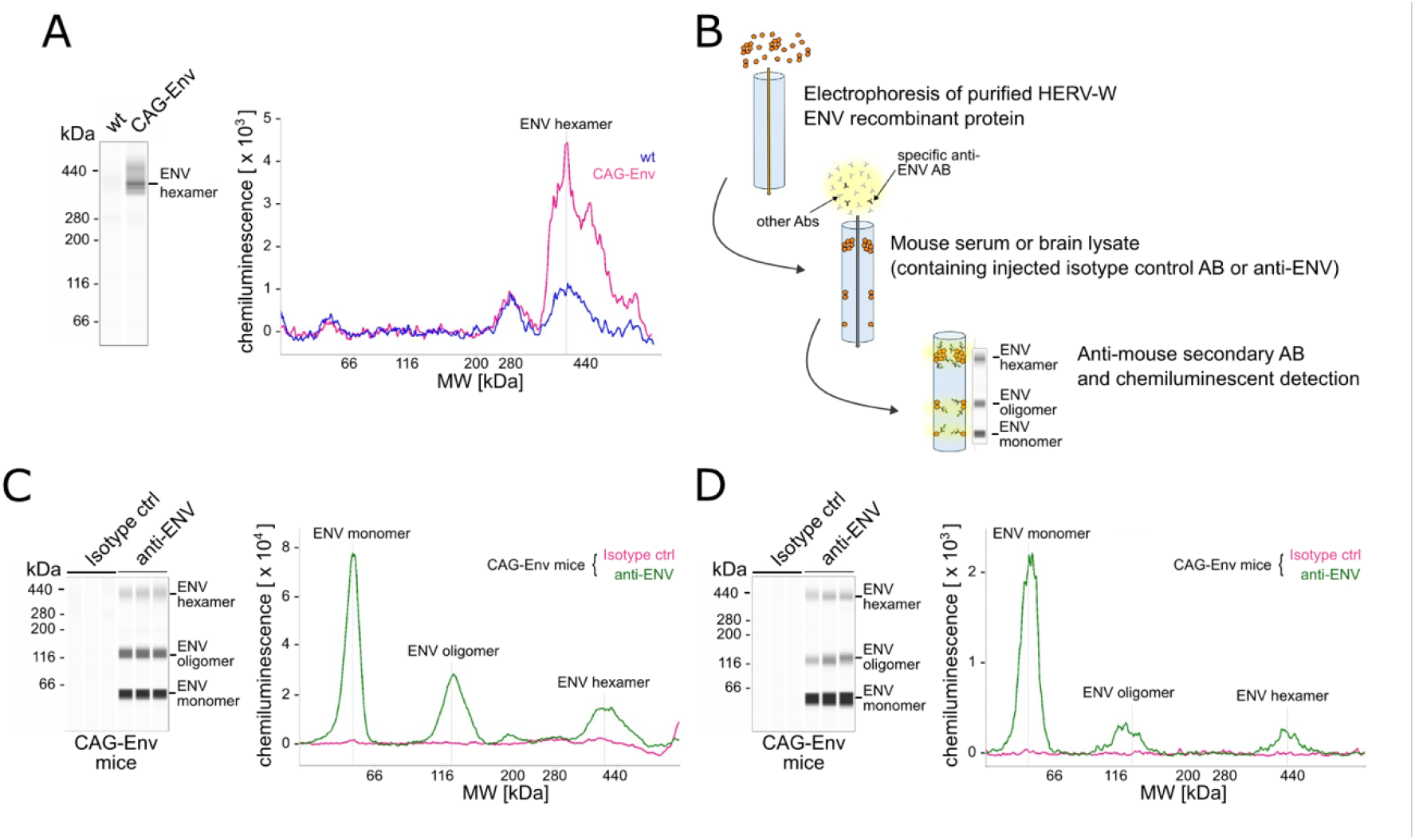
Anti-ENV antibodies can cross the blood-brain barrier when injected in the periphery. (A) Representative detection of hexameric HERV-W ENV protein in CAG-Env transgenic (CAG-ENV, pink) but not in wildtype (wt, blue) litter-mate mice. (B) Schematic presentation of anti-ENV antibody detection in non-fixed brain tissue or blood serum, also referred to as tissue serology. (C) Anti-ENV antibody detection in sera of CAG-Env transgenic mice previously receiving two anti-ENV injections (i.p., green) but not in isotype control antibody-injected mice (pink) after 7 weeks of cuprizone-induced demyelination. (D) Anti-ENV antibody-injected-(green) and control antibody-administered-(pink) transgenic mice were used to prepare PBS-perfused, non-fixed brains and to detect the peripherally applied (i.p.) anti-ENV antibody in the brain parenchyma. Data derived from n=1 (A) and n=3 (C,D) mice, respectively.

### Remyelination-related oligodendroglial parameters

Rescued myelinated areas with the corpus callosum of transgenic mice treated by the anti-ENV antibody (Fig. 1B,D) can either result from prevented demyelination and/or fostered myelin repair activities. We therefore determined the expression of several oligodendroglial proteins known to mark and regulate differentiation of resident oligodendroglial precursor cells (OPCs). The analysis was conducted at two time points during demyelination (5 wk and 7 wk CPZ) and after two weeks of remyelination (2wk rem; see Fig. 1A for presentation of the experimental flow). Immunofluorescent histology revealed increased densities of platelet-derived growth factor receptor-α (Pdgfrα)-positive cells, representing the resident OPC pool (Fig. 3A,B), which also appeared to proliferate more in presence of the anti-ENV antibody (Ki67-positivity, Fig. 3A,C). Subsequently, SRY-Box transcription factor 10 (Sox10) and adenomatous-polyposis-coli (Apc) staining was conducted. Over the entire investigation period, an increased density of Sox10-positive cells was observed (Fig. 3A,D). Sox10-positive/Apc-negative cells correspond to differentiating oligodendroglia and were found to be significantly elevated in mice injected with the anti-ENV antibody as compared to mice receiving the control antibody (Fig. 3A,E). The number of Sox10/Apc double-positive maturing oligodendrocytes was also found to be elevated in transgenic brains, particularly during the remyelination phase (Fig. 3A,F). Additionally, the densities of early (re-)myelinating oligodendrocytes, identified by their expression of the breast carcinoma amplified sequence 1 (Bcas1) protein, were significantly elevated upon anti-ENV treatment at the conclusion of demyelination and during the subsequent remyelination process (Fig. 3A,G). These observations demonstrate that the application of the anti-ENV antibody positively impacts oligodendroglial differentiation *in vivo*, resulting in enhanced remyelination.

**Figure 3:**
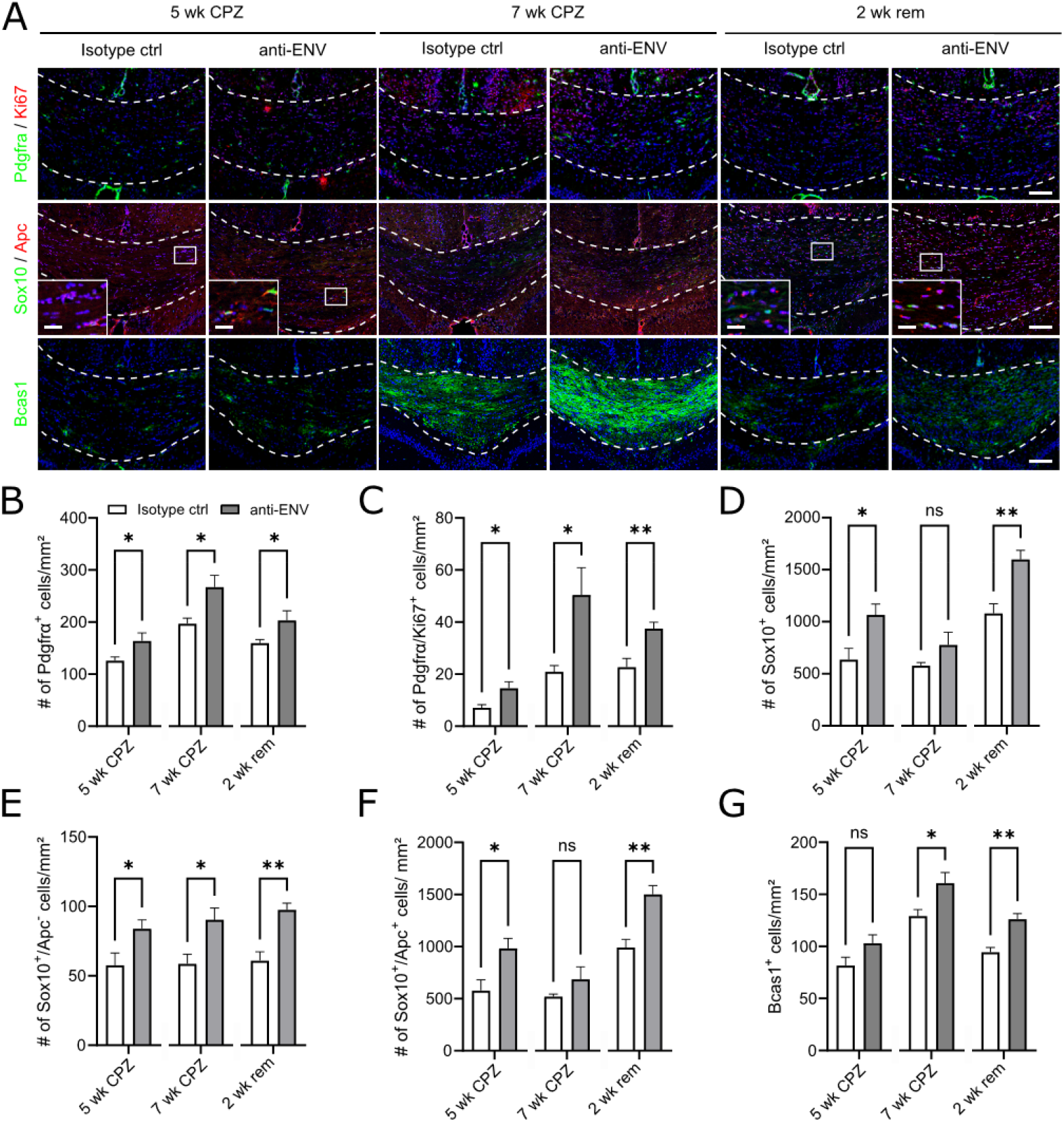
Anti-ENV application fosters oligodendroglial differentiation, proliferation, and maturation towards myelinating cells. (A) Representative immunohistochemical images of Pdgfrα/Ki67-, Sox10/APC-, and Bcas1-expression in transgenic (CAG-Env) corpora callosa after 5 and 7 weeks of CPZ-administration (5 wk CPZ, 7 wk CPZ) and after 2 weeks of CPZ withdrawal (2 wk rem) upon isotope control antibody or anti-ENV antibody injections. (B) Quantification of Pdgfrα-positive-, (C) Pdgfrα/Ki67 double-positive-, (D) Sox10-positive-, (E) Sox10-positive/Apc-negative-, (F) Sox10/Apc double-positive-, as well as of (G) Bcas1-positive cell densities of control antibody-injected vs. anti-ENV-injected transgenic mice. Data are presented as mean values (n=6) ± SEM. Significance was analyzed by Student’s two-sided, unpaired t-test and Mann-Whitney *U* two-tailed test (95% CI) at *P < 0.05, **P < 0.01, ns = not significant. Dashed lines in (A) indicate the area of corpus callosum. (Scale bars: 100 µm and for blow-ups 25 µm).

### Reduced generation of neurotoxic microglia and astroglia

We previously reported that the HERV-W ENV protein polarizes microglial cells *ex vivo* (human and rat primary cells) and *in vivo* (transgenic mouse model) towards an axon-damaging phenotype (13, 14). Furthermore, a transcriptomic analysis of microglial cells challenged with ENV protein further corroborated the presence of reactive, neurotoxic cells (20). Thus, the expression of several microglial proteins known to mark distinct polarisation states was assessed histologically in response to anti-ENV antibody treatment at three different time points during corpus callosum demyelination and remyelination. By analyzing the degree of ionized calcium-binding adapter molecule 1 (Iba1)-positive areas, a significant reduction in the density of microglial cells was detected in anti-ENV-treated animals after 7 weeks (end of demyelination; Fig. 4A,B). The use of additional markers, such as C-type lectin domain family 7 member A (Clec7a), which is induced and associated with microglia in neurodegenerative diseases, or transmembrane protein 119 (Tmem119), a protein marking homeostatic microglial cells, further corroborated these findings. Homeostatic cells (Tmem119-positive, Clec7a-negative) were gradually elevated in anti-ENV-treated mice (as opposed to control antibody-treated animals) reaching a significant increase during remyelination (Fig. 4A,C). On the other hand, neurotoxic cells (Tmem19-negative, Clec7a-positive) were significantly reduced at the end of the demyelination period when the ENV protein was neutralized by the antibody (Fig. 4A,D). Notably, Tmem119/Clec7a-double-positive intermediate cell stages also increased significantly during remyelination (Fig. 4A,E). This suggests that the parenchymal presence of the anti-ENV antibody and its subsequent neutralization of the HERV-W ENV protein permitted a phenotypic transformation within the microglial cell population.

**Figure 4:**
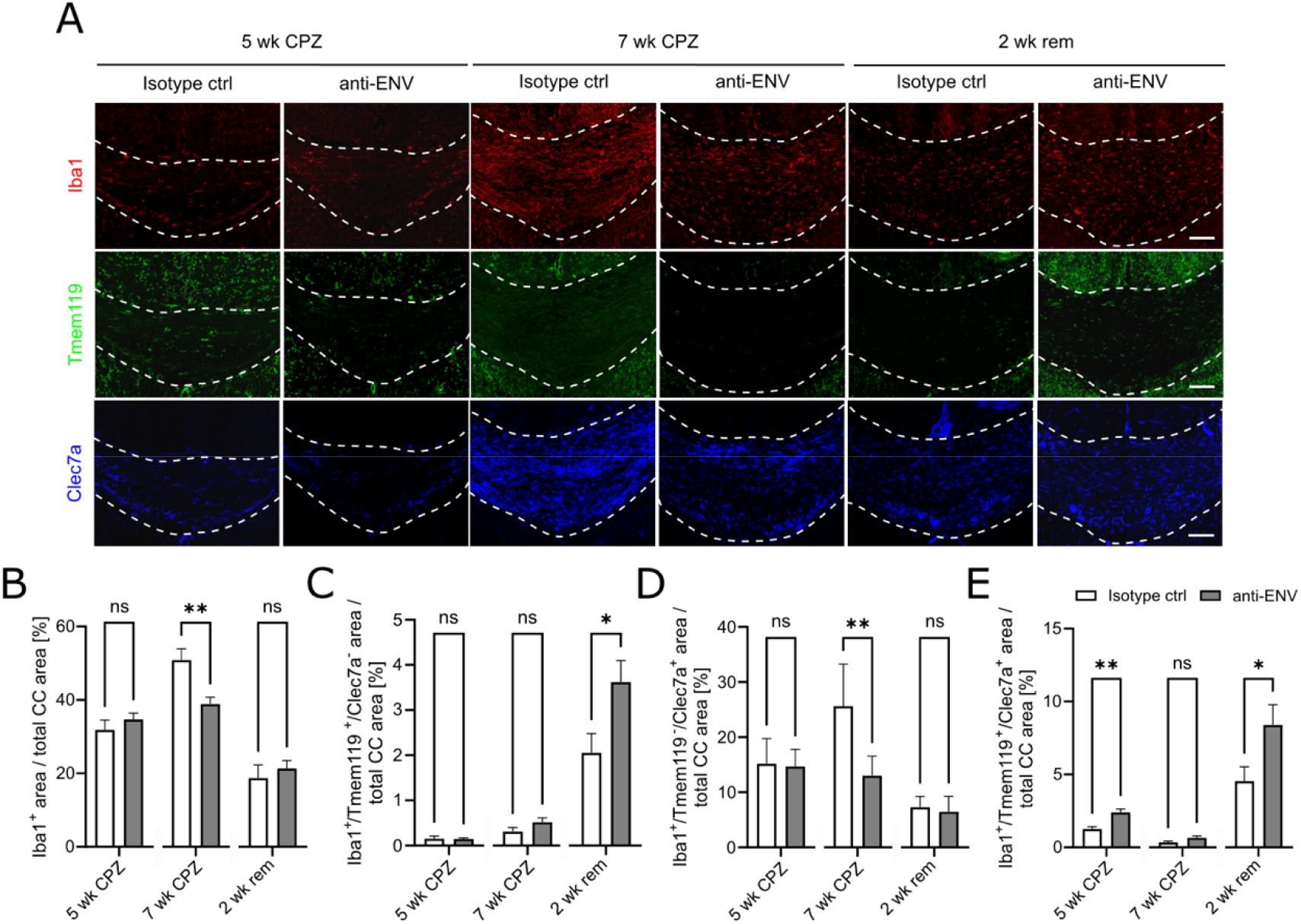
Anti-ENV application promotes homeostatic microglial polarization while neurotoxic microglial traits are reduced in transgenic CAG-Env mice. (A) Representative immunohistochemical pictures of Iba1-, Tmem119- and Clec7a-expressing cells in transgenic corpus callosum (CC) tissues (control antibody and anti-ENV-treated animals). (B) Quantification of the Iba1-positive-, (C) Iba1-positive/Tmem119-positive/Clec7a-negative-, (D) Iba1-positive/Clec7a-positive/Tmem119-negative-, as well as of (E) Iba1/Tmem119/Clec7a triple-positive areas over the total corpus callosum area in isotype control antibody-vs. anti-ENV injected transgenic mice at 5 and 7 weeks of CPZ-treatment and after 2 weeks of remyelination. Data are presented as mean values (n=6) ± SEM. Statistical significance of histological data was analyzed via Student’s two-sided, unpaired t-test and Mann-Whitney *U* two-tailed test. Data were considered statistically significant (95% CI) at *P < 0.05, **P < 0.01, ns = not significant. Dashed lines demarcate the corpus callosum. (Scale bars: 100 µm).

In addition to microglia, our previous study on HERV-W ENV’s *in vivo* effects also provided strong evidence of a pathological involvement of astroglial cells (14). Specifically, we observed not only direct ENV-mediated effects on astrocytes, but also indirect responses induced by ENV-activated microglia, which likely contribute to the generation of reactive, neurotoxic astrocytes (20). Thus, the expression of several astrocytic proteins known to mark different pathological states was assessed histologically at the same time points. The overall glutamate aspartate transporter 1 (Glast1)-positive area remained unchanged between the two different antibody treatments at all three time-points (Fig. 5A,B). However, using S100a10, an anti-inflammatory and restorative astroglial marker (21), or lipocalin2 (Lcn2), a previously assigned neurotoxic astrocyte marker (14), we identified astroglial changes in response to ENV neutralization. Glast1-positive cells expressing the S110a10 protein but with absent Lcn2 signals were boosted upon anti-ENV antibody treatment at late demyelination and in the remyelination phase (Fig. 5A,C). Neurotoxic astrocytes (lacking S100a10 but expressing Lcn2) reacted inversely and were reduced at all three time-points reaching a significant reduction during acute demyelination and at remyelination (Fig. 5A,D). Notably, S100a10/Lcn2-double-positive intermediates were also significantly diminished during remyelination (Fig. 5A,E). Overall, the observed astrocytic responses were highly reminiscent of microglial reactions (Fig. 4), corroborating a combined effect of the anti-ENV antibody in re-establishing a less hostile parenchymal glial environment.

**Figure 5:**
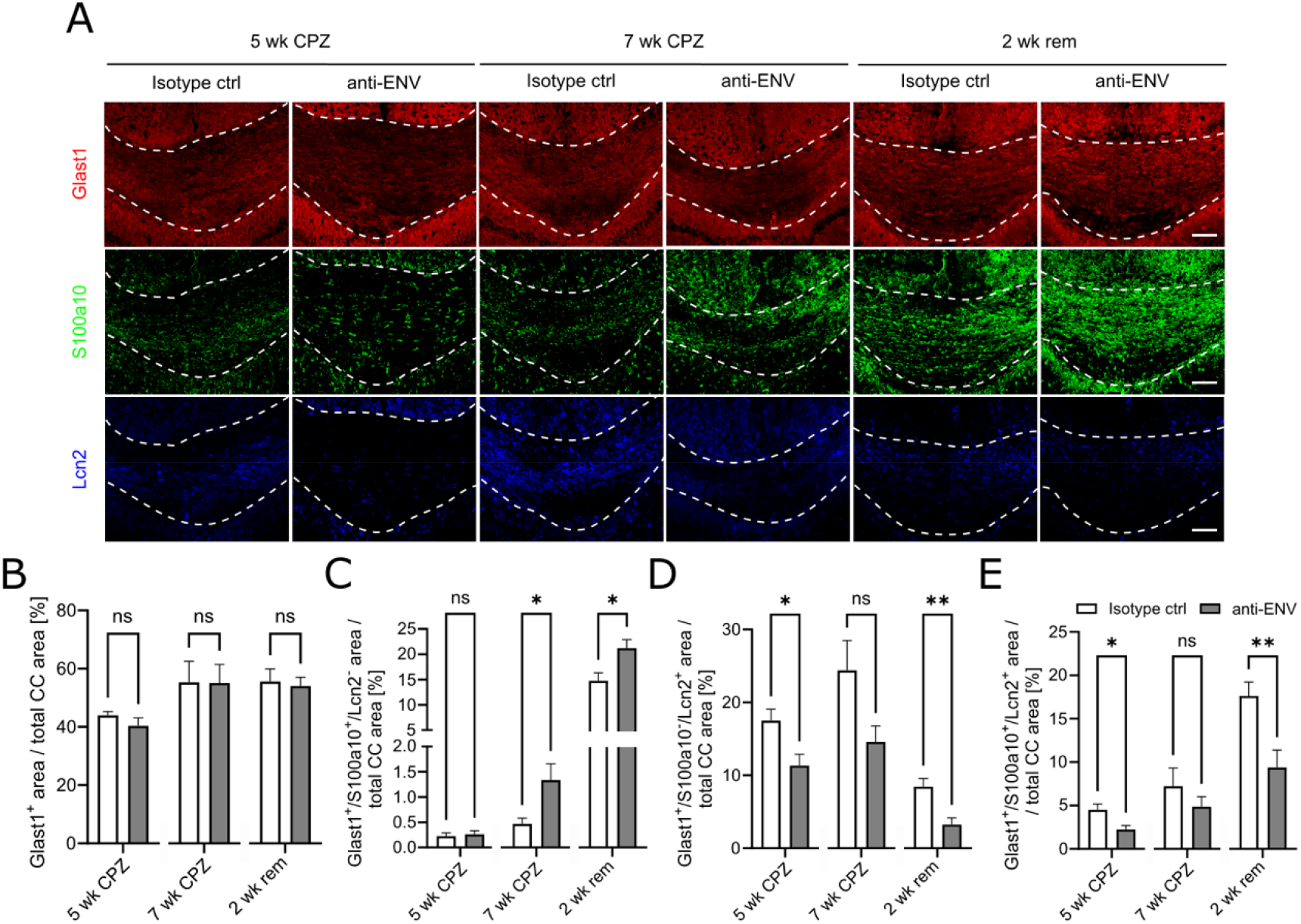
Anti-ENV injection fosters a restorative astroglial cell response in transgenic CAG-ENV mice. (A) Representative immunohistochemical images of Glast1-, S100a10- and Lcn2-expressing cells in transgenic corpus callosum tissue sections administered upon isotope control antibody or anti-ENV antibody injection. (B) Quantification of Glast1-positive areas within the corpus callosum of CAG-Env mice (control- and anti-ENV-injected). (C) Quantification of Glast1/S100a10 double-positive/Lcn2-negative-, (D) Glast1/Lcn2-double-positive/S100a10-negative-, and (E) Glast1/S100a10/Lcn2 triple-positive areas within the corpus callosum of transgenic mice (control- and anti-ENV-injected). Data are presented as mean values (n=6) ± SEM. Significance was analyzed via Student’s two-sided, unpaired t-test and Mann-Whitney U two-tailed test (95% CI) at *P < 0.05, **P < 0.01, ns = not significant. CC = corpus callosum. Dashed lines indicate the area of the corpus callosum. (Scale bars: 100 µm.)

### Peripheral biomarker detection

HERV-W is a human-specific pathological entity, and its envelope protein is therefore considered to exert pathological functions that can only be mimicked to a limited extent by animal models. Consequently, important functional insights into its role in MS have only been gained from the recently published investigation of the engineered transgenic mouse line (14) (Gruchot et al., 2023) in combination with CPZ and EAE models. The subsequent transcriptomic description of cellular responses established that HERV-W is primarily involved in PIRA-related smouldering diseases processes (17, 20). To further connect the currently existing *ex vivo* and *in vivo* data gathered from ENV challenges on isolated human cells and in animal models with findings from patients treated with the neutralizing anti-ENV antibody Temelimab (15); Piehl et al., ECTRIMS 2024, full manuscript in preparation), we conducted peripheral biomarker testing in our animal model. Although an extended knowledge on suitable biomarkers for disease progression (in both animals as well as patients) is still developing, serum released neurofilament light chain (sNfL) has been recognized as a reliable peripheral marker for MS-related neuron/axon degeneration, flagging a higher risk of disability progression, eminent and depicting PIRA-related processes (22).

We determined sNfL concentrations in two distinct experiments. In the initial setup, we compared its concentration in wildtype littermates vs. ENV-expressing transgenic mice at the conclusion of CPZ-dependent demyelination (time point 7 weeks). Here, we detected a significant increase in sNfL levels in the transgenic background (Fig. 6A) consistent with the observed elevated APP spheroids as signs of increased neurodegeneration (14). A second determination of sNfL levels was subsequently conducted using ENV transgenic mice exclusively, but this time, they were either treated with the isotype control antibody or the anti-ENV antibody (two injections) at the time point 7 weeks of CPZ administration. Here, a notable decrease in sNfL levels was observed in transgenic mice receiving anti-ENV antibody (Fig. 6B), in line with the diminished detection of APP spheroids (Fig. 1C,D). Higher sNfL levels in response to transgenic ENV expression further support its harmful effects *in vivo*. Our findings also allow us to compare it directly with patient data, where sNfL levels were similarly reduced in response to Temelimab treatment (Piehl et al., ECTRIMS 2024, full manuscript in preparation). It is important to note that in this phase 2a clinical trial (NCT04480307), Temelimab was administered in combination with Rituximab, thereby enabling evaluation of mainly progression-related processes.

**Figure 6:**
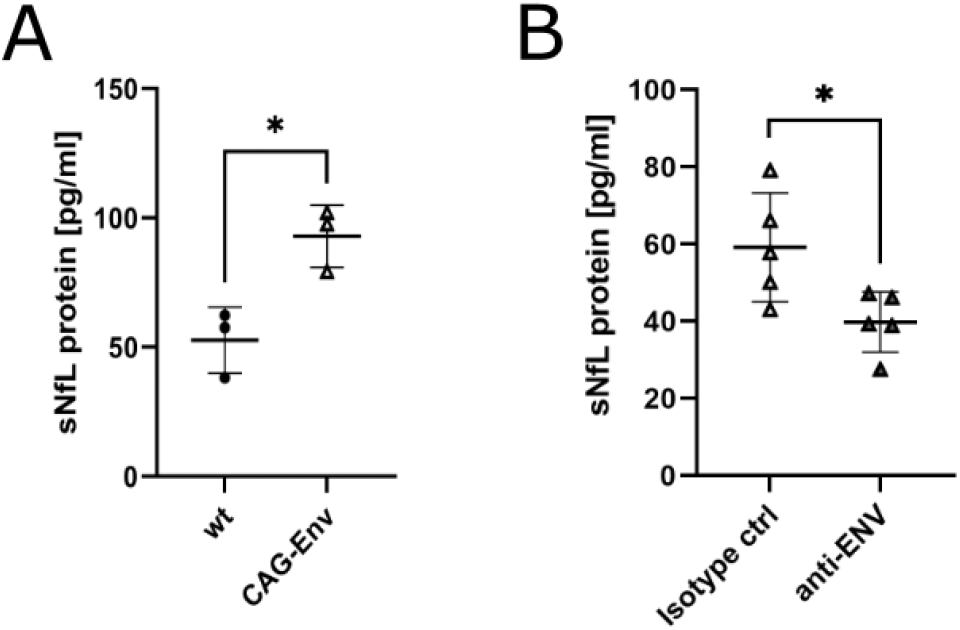
Serum neurofilament light chain (sNfL) profiles in mouse sera. (A) Serum protein detection in wildtype versus CAG-Env transgenic revealed a significant increase in sNfL levels in presence of ENV protein after 7 weeks of CPZ-treatment. (B) Anti-ENV antibody treatment of ENV expressing transgenic mice after 7 weeks of CPZ-dependent demyelination and two antibody injections significantly diminished sNfL levels compared to isotype control antibody administration. Data are presented as mean values deriving of n=3 (A) and n=5 (B) ± SEM. Significance was assessed by Student’s unpaired t test. Data were considered as statistically significant (95% CI) at *P < 0.05.

## Discussion

Prior to our recent investigation using a genetically engineered transgenic mouse model that mimics the expression of the HERV-W ENV as observed in people with MS (14), functional assessment of HERV-W was restricted to inferred processes derived from histology in conjunction with *ex vivo* functional assays conducted with primary cells (12, 13). Nevertheless, this provided evidence that oligodendroglia and microglia were among the most strongly affected cell types under conditions of HERV-W ENV expression.

Using a mouse model recapitulating human-specific expression of the HERV-W ENV, we identified marked pathological reactions of all three glial cell types to the presence of this endogenous retroviral element, particularly under MS-related conditions, demonstrating a neurotoxic environment and impaired myelin regeneration in presence of the ENV protein. Notably, axon degeneration and destruction were also observed, even in a mild CPZ model, as well as in EAE mice (14). This suggests that HERV-W promotes disease progression and predominantly impacts PIRA-related processes. This observation is consistent with imaging results from anti-HERV-W (Temelimab)-treated MS patients (15)). Considering these studies, it was therefore tempting to study the potential effects of an ENV-directed and neutralizing antibody in transgenic mice.

*In vivo* experimental data presented here provide critical new evidence that intraperitoneally injected anti-ENV (the murine parental antibody of the humanized Temelimab developed for clinical use) indeed accumulated in the parenchyma and exhibited ENV binding activity, thereby recognizing all previously described and pathologically relevant ENV forms such as hexamers, oligomers, and monomers (19). Given that the CPZ demyelination model was chosen to focus on neural cell responses, the blood-brain barrier (BBB) was expected to remain largely intact. This suggests that the overall antibody penetration is sufficiently high to effectively neutralize the transgenically expressed ENV protein. As a result, enhanced remyelination was observed following anti-ENV treatment, as OPCs were more proliferating and were also rescued in their differentiation process. Additionally, it was observed that depolarizing microglia and astroglia resulted in the induction of more favorable, less hostile, and regenerative phenotypes. Moreover, the presence of the anti-ENV antibody significantly reduced APP-spheroid generation (14, 18), indicating preservation of axons. Axonal protection on the other hand is the most likely underlying cause of reduced atrophy rates and diminished black hole appearance in Temelimab-treated patients (15), additionally complementing our current view of HERV-W activity in the autoimmune brain.

The primary finding of this study is therefore that neutralizing the activity of the CNS-expressed and resident ENV protein through a specific antibody in the periphery results in the transfer and accumulation of the antibody in the CNS parenchyma. Furthermore, this approach targets all glial cell types previously demonstrated to respond to the ENV protein directly and hence addresses mainly PIRA-related pathological processes. This is of importance as currently, there is no drug to treat PIRA in a clinically meaningful extent. Ocrelizumab, the only licensed drug to treat primary progressive (PP)MS shows, based on its mode of action, minimal effect of disability worsening by suppressing subclinical inflammatory effects, while there is accumulating evidence of a lack of effect progression in its strict sense (23). Hence, novel approaches are needed to address the unmet medical need for therapies specifically tackling PIRA.

In general, there is an ultimate need for drugs designed to target pathological processes behind the blood-brain-barrier and in particular to modulate the activity of parenchyma-resident cells. This is a particularly challenging task when using antibodies which have also already been explored in the context of myelin repair. Opicinumab, an anti-LINGO-1 antibody designed to foster remyelination in multiple sclerosis and optic neuritis (24, 25) has been investigated (26). These studies highlighted limitations not only in clinical treatments aimed at facilitating myelin repair, readouts, and endpoints but also in reaching targets with pharmacologically relevant doses. Nevertheless, such therapeutic approaches should also be considered to ameliorate the pathogenic effects of microglia and astrocytes, potentially involving the identification of suitable small molecules (*de novo* or repurposed) and the application of antibodies.

Our initial findings in the relatively mild CPZ model, which usually shows minimal axon degeneration (27), revealed that the ENV transgenic background significantly increased local axon destruction. A similar elevation was also observed in EAE (14). When translated to people with MS this observation may be particularly relevant in the context of PIRA, where mechanisms other than mediated by infiltrated immune cells must account for tissue destruction. Consequently, it was suggested that HERV-W ENV primarily influences smouldering neuroinflammatory processes (17). This hypothesis is now further supported by the findings presented here. Applying the anti-ENV antibody significantly reduces this crucial aspect of tissue destruction and irreversible functional loss. This was accompanied by a diminished NfL release in the mouse serum. These findings align well with previous research showing a reduction in atrophy and black hole formation in Temelimab-treated multiple sclerosis patients (15) and a decrease in sNfL levels in B cell-depleted patients upon Temelimab co-administration (Piehl et al., ECTRIMS 2024, full manuscript in preparation).

sNfL and sGFAP serve as tissue damage and sclerosis markers, respectively, reflecting disease activity and heralding disease progression. In both the mouse model used here and in people with MS, the presence of NfL in serum likely reflects acute neuroaxonal injury (28, 29). The observed changes of sNfL appearance are therefore congruent between mouse model and Temelimab-treated human patients. Whereas in the Temelimab-treated patient cohort, peripheral sGFAP levels as a measure of astrocytosis and scar formation also dropped (Piehl et al., ECTRIMS 2024, full manuscript in preparation), sGFAP was not measurable in the transgenic mouse model for technical reasons – a limitation that shall be addressed in follow-up studies.

As the focus of the anti-ENV antibody application was on neural cell fates and tissue recovery rates, best mimicking relapse-free conditions, we explicitly excluded any investigations on lymphocyte-directed effects. This, even though in the EAE model, ENV transgene expression led to disease exacerbation, increased infiltration of CD3-positive T cells, larger white matter lesions, and a degree of disease chronification (14). Notably, the CHANGE-MS clinical trial did so far not support a direct effect of HERV-W ENV on lymphoid cells (15). While a more detailed description of immune cells in presence of the ENV protein and upon peripheral anti-ENV application is required to fully understand the contribution of HERV-W activation to disease evolution, this was beyond the scope of this study and will be conducted in subsequent investigations. These will also include analyses of stem cell niches, endothelia and pericytes, all of which are functionally implicated in autoimmune and neurodegenerative pathologies.

## Materials and Methods

### Ethics statements for animal experiments

All animal experiments were conducted in compliance with the ARRIVE guidelines and the National Institutes of Health guide for the care and use of Laboratory animals (NIH Publications No. 8023, revised 1978). Experimental procedures involving animals were approved by the Institutional Review Board (IRB) of the ZETT (Zentrale Einrichtung für Tierforschung und wissenschaftliche Tierschutzaufgaben) at the Heinrich-Heine-University Düsseldorf under internal licenses O69/11, O90/15. Additional approvals for animal procedures were obtained from the review board of the state government LANUV (Landesamt für Natur, Umwelt und Verbraucherschutz Nordrhein-Westfalen, North-Rhine Westphalia, Germany) under license Az 81-02.04.2019.A203. All procedures had received prior approval from the appropriate veterinary authority.

### Animal models

We used transgenic C57BL6/J;129P2/Ola-Hprttm(CAG-Env) (CAG-Env; 75% C57BL6/J and 25% 129P2/Ola) mice. Genotypes were determined by genomic PCR targeting hypoxanthine-guanine phosphoribosyltransferase (Hprt) alleles, as described previously (14). Mice were bred and kept in the ZETT at Heinrich-Heine-University Düsseldorf under strict pathogen-free conditions, with a 12-hour light/dark cycle and unlimited access to food and water. Corpus callosum demyelination was initiated in 8-week-old mice by feeding a diet containing 0.2% (w/w) cuprizone [bis(cyclohexanone)oxaldihydrazone] (V-1534, Ssniff, Soest, Germany) as previously described (14, 21, 30, 31). To achieve consistent demyelination, animals were administered cuprizone (CPZ) for 7 weeks (Fig. 1A) followed by recovery on normal chow (V-1534, Ssniff) for 2 weeks (2-week remyelination). Anti-ENV01 antibody (GeNeuro, Geneva, Switzerland) or isotype control antibody (InVivoMAb mouse IgG1 MOPC-21, Biozol) injections (i.p., 30 mg/kg) started after 3 weeks of CPZ-treatment followed by a new injection every second week resulting in up to 3 total injections (Fig. 1A). Diets were replaced twice weekly, and body weight was monitored twice per week. Each experimental group consisted of six animals (either sex) per time point, as determined by cohort size analysis using G*Power 3.1.9.7 (effect size: 2.6; α = 0.05; power = 0.95).

### Tissue isolation for protein expression analysis and immunohistology

Briefly, animals were deeply anesthetized with isoflurane and transcardially perfused with 20 mL cold phosphate-buffered saline (PBS) to remove blood cells from the brain tissue. For the detection of HERV-W ENV protein as well as of injected antibodies using automated western blot techniques, whole brains were isolated and immediately frozen in liquid nitrogen. All samples were stored at -80°C until protein isolation. For immunohistology of CPZ treated mice brains were fixed and processed as described previously (14).

### Simple Western analysis for HERV-W ENV protein and antibody detection

Brain tissues were extracted in complete permeabilization buffer (Thermo Fisher Scientific, USA) and incubated on ice for 10 minutes. After centrifugation at 16,000 g for 15 minutes, the supernatant was then subjected to the Simple Western system (Bio-Techne, USA) based on capillary electrophoresis (CE) which offers a size assay combining CE-SDS with immunodetection to separate proteins by molecular weight. Sample loading and detection were automated, fast and quantifiable. Transgenic HERV-W-ENV protein was detected with biotinylated Temelimab primary antibody (Geneuro) and a secondary streptavidin detection module (DM-004, Bio-Techne) as previously described in (19). Quantification of chemiluminescence was based on peak height after correction for a baseline signal. Raw data were generated by Compass software (Bio-Techne). Detection of the ENV01 antibody in mouse sera and brains was also performed using Simple Western analysis. In this serological format, recombinant HERV-W ENV protein (GTP Technology, France) was used as the antigen, while sera and brain extracts were used as primary antibody sources. Tissues were extracted in RIPA buffer (Sigma Aldrich, USA) with antiprotease inhibitors cocktail and supernatants were kept after centrifugation at 10,000g at 4°C. A secondary mouse antibody module was used to complete the immunoassay (Bio-Techne, ref. DM-002). Quantification of chemiluminescence was based on peak height after correction for a baseline signal. Raw data were generated by Compass software (Bio-Techne, USA).

### Immunohistochemical procedures

Tissue isolation, fixation, and coronal sectioning of the caudal corpus callosum (Bregma -0.70-2.06; 12 µm) were prepared and preserved according to (14). To evaluate the relative myelination of the corpus callosum, Luxol fast blue (LFB, Sigma-Aldrich) staining was performed as previously outlined (Gruchot et al., 2023). For immunohistochemistry, brain sections were thawed, dried for 15□min at RT, then rehydrated for 5 min in distilled water, further post fixated for 5 min in 4% PFA and finally for another 5 min in -20°C acetone. Sections were washed once in Tris-buffered saline (TBS, pH 7.6), once in TBS-T (TBS containing 0.02% Triton) for 5 min each and incubated for another 5 min in 0.3% H2O2 solution. Blocking was performed using 10% NGS (Sigma-Aldrich) and 5% biotin-free bovine serum albumin (BSA; Sigma-Aldrich; in TBS-T) for 30□min at RT, followed by the application of rabbit anti-amyloid precursor protein (APP; 1/200; Thermo Fisher Scientific, RRID:AB_2533902) in 10% NGS and 5% BSA in TBS over night at 4 °C. Afterwards, sections were washed twice in TBS (5 min each) and a biotinylated secondary antibody (goat anti-rabbit [1/200; Vector Laboratories, Burlingame, CA, USA]) was added for 30 min. Next, sections were washed twice in TBS and ABC reagent was incubated for another 30 min according to the manufacturer’s protocol (Vectastain Elite ABC HRP kit; Vector Laboratories). Afterwards, sections were washed again twice for 5 min in TBS and peroxidase substrate was added for 5 min at RT (ImmPact DAB; Vector Laboratories). The reaction was stopped by two washing steps in ddH2O, followed by hematoxylin nuclear stain (Carl Roth), dehydration and embedding in ROTI-Histokitt II (Carl Roth). For immunofluorescence staining, brain sections were thawed, rehydrated for 5 min in distilled water, post fixated for 5 min in 4% PFA and for another 5 min in -20°C acetone. Before blocking, sections were washed once using Tris-buffered saline (TBS, pH 7.6) and once in TBS-T (TBS containing 0.02% Triton) for 5 min each. Blocking was performed with 10% serum according to the host of the secondary antibody (NGS or NDS respectively; Sigma-Aldrich) and 5% biotin-free BSA (Sigma-Aldrich; in TBS-T) for 30□min at RT, followed by application of the following antibodies (in 10% NGS/NDS, 5% BSA, TBS) and incubation overnight: goat anti-Pdgfrα (1/250; R and D Systems RRID:AB_2236897), rabbit anti-Ki67 (1/250; Abcam; RRID:AB_302459) rabbit anti-Sox10 (1/100, DCS Immunoline, Hamburg, Germany; RRID: AB_2313583), mouse anti-APC (CC1, 1/300, Sigma-Aldrich; RRID:AB_ 2057371), mouse anti-Bcas1 (1/200; Santa Cruz; Dallas, TX, USA; RRID:AB_10839529), rabbit anti-Iba1 (1/500, WAKO Pure Chemical Corporation; RRID: AB_839504), rat anti-Clec7a (Dectin1; 1/50; InvivoGen San Diego, CA, USA; RRID:AB_2753143), anti-S100a10 (1/500; Thermo Fisher Scientific; RRID: AB_2092361) chicken anti-Iba1 (1/500; Aves; RRID:AB_2910556), rabbit anti-Tmem119 (1/100; Abcam; RRID:AB_2921338), goat anti-Lcn2 (1/100; R and D Systems; RRID:AB_355022) and mouse anti-Glast1 (1/600; Miltenyi, RRID:AB_10829302). Sections were washed two times for 5 min in TBS and incubated with the species-appropriate Alexa fluorochrome-conjugated secondary antibody (1/200 in TBS, Thermo Fisher Scientific) and 4′, 6-diamidino-2-phenylindole (DAPI; 20 ng/ml, Roche) for 30□min at RT. Afterwards, sections were washed once in TBS and once in TBS for 5 min each and embedded using Shandon™ Immu-Mount (Thermo Fisher Scientific).

### Serum protein determination

Levels of serum NfL were quantified using the fully automated immunoassay platform, Ella (Bio-techne, USA). All proteins were quantified using a single disposable microfluidic SimplePlex^™^ cartridge. Serum samples were thawed on ice and diluted 1:2 in sample diluent and loaded into cartridges (ref SPCKB-MP-003168, USA) with relevant high and low control concentrates. All steps in the procedure were run automatically by the instrument. The obtained data were displayed as pg/mL and automatically calculated by the internal instrument software (Simple Plex Explorer, Bio-Techne, USA).

### Image acquisition and analysis

Images of LFB and DAB-stained tissue sections were captured on an Axioplan 2 microscope (Zeiss, Jena, Germany), while the other images were taken at a Zeiss CLSM microscope 510 (CLSM 510, Zeiss) strictly using the same exposure times, laser intensities and digital gains. The quantification was performed using ImageJ software (National Institute of Health (NIH) Bethesda, MD, USA) analyzing 3 sections (per condition/replicate) for each marker setup as previously described in Gruchot et al., 2023. Analysis of the APP-positive spheroids were quantified manually, using the ImageJ tool “cell-counter”. To assess the relative myelination of the corpus callosum, images of LFB staining were transformed to grey-scale, and the same threshold was applied to all images to create binary images. Afterwards the area of the corpus callosum as well as the LFB-positive area was assessed to calculate the relative LFB-positive myelinated areas. The analysis of double- and/or triple immune-positive cells was conducted as previously described (14) using ImageJ software.

### Statistical analysis

Data are presented as mean values ± standard error of the mean (SEM) in which n represents the number of independent replicates. Statistical analyses were conducted using Graph-Pad Prism 8.4.3 (GraphPad Software, San Diego, California USA). Shapiro-Wilk normality test was used to assess the absence of Gaussian distribution of all datasets. To determine statistical significance for normally distributed data sets, the Students t-test was applied while the Mann– Whitney U test was used when not passing the Shapiro–Wilk normality test. The experimental groups were considered significantly different at *p<0.05, **p<0.01, ***p<0.001.

## Data and materials availability

All data generated or analyzed during this study are included in this published article.

## Acknowledgments

We acknowledge Zippora Kohne, Birgit Blomenkamp-Radermacher, Brigida Ziegler for their technical assistance.

